# Laminin-Derived Peptides Bind Neuropilin-1 at the VEGF-A_165_ Binding Pocket

**DOI:** 10.64898/2026.10.27.762289

**Authors:** Evan Veltri, Sándor Lovas

**Affiliations:** Department of Biomedical Sciences, Creighton University, Omaha, Nebraska 68178, United States

## Abstract

Neuropilin-1 (NRP-1) is associated with glioblastoma progression and contains a b1-do-main pocket that binds the C-terminal region of VEGF-A165. Whether laminin-derived peptide fragments can associate with this site is unknown. AlphaFold 3 complex prediction followed by 15 μs of all-atom production molecular dynamics simulation, trajectory-based pocket-occupancy and interaction analyses, and FoldX energetic scoring was performed to compare binding of the laminin-derived pentapeptide Ac-Tyr-Ile-Gly-Ser-Arg-NH_2_ (peptide **1**)with its retro (Ac-Arg-Ser-Gly-Ile-Tyr-NH_2_, peptide **2**) and retro-inverso (Ac-arg-ser-Gly-ile-tyr-NH_2_, peptide **3**) analogs. Peptide **1** showed the highest mean equilibrated occupancy of the canonical VEGF-A165 pocket (60.3 ± 34.6%) compared with peptide **2** (25.9 ± 32.7%) and peptide **3** (21.4 ± 29.5%), while sampling interactions with canonical pocket residues including Asp320, Tyr297, and Tyr353; FoldX binding-energy estimates were similar across peptides despite their differing pocket persistence. Complementary nanoDSF measurements using recombinant NRP-1 and synthetic peptides yielded apparent dissociation constants in the low-micromolar range for all three peptides (12.7 µM – 33.3 µM), providing experimental support for the predicted interactions. These results support persistent, sequence-dependent association of peptide **1** with the VEGF-A_165_ binding site of b1-NRP-1 and identify peptide analogs as a candidate for further optimization as an NRP-1-binding peptide.

## 2. Introduction

Glioblastoma (GB) is the most common primary malignant brain tumor and one of the most aggressive neoplasms found in the central nervous system^1–2^. GB is rapidly progressing and highly invasive with a five-year survival rate of less than 7% and a median overall survival of one to two years^1–3^. Approximately 52.2% of malignant primary brain and 13.7% of all primary CNS tumors in the United States is typically diagnosed in older adults^4^. Global incidence ranges from 0.59 to 5 per 100,000 people every year, with trends suggesting incidence is on the rise^2^. Even with maximal safe surgical resection followed by radiotherapy and chemotherapy via the DNA-alkylating agent temozolomide (TMZ), the recurrence is quite universal and fatal^1,3,5^. Patients who have a O6-methylguanine-DNA methyltransferase (MGMT) promoter methylation have improved results on TMZ, and MGMT promoter methylation predicts improved temozolomide response and better survival, although outcomes remain poor even in this subgroup^6^. The dire combination of poor prognosis, aggressiveness, and limited therapeutic options highlights the urgent need for new therapeutic strategies for GB.

GB is known for its extensive neovascularization and highly invasive phenotype, characteristics that are bolstered by its abnormal expression of cell-surface receptors that regulate angiogenesis, immune evasion, and cell migration^7–9^. Among these cell-surface receptors, neuropilin-1 (NRP-1) has been identified as an important mediator of GB progression^7–9^. NRP-1 is a single-pass transmembrane glycoprotein that binds a variety of extracellular ligands. It has five extra-cellular domains (a1, a2, b1, b2, and c) and a short intracellular GAIP-interacting protein (GIPC) region^7–8^. NRP-1 is significantly overexpressed in GB and contributes to its dismal outcomes by enhancing angiogenesis, facilitating interactions with the extracellular matrix (ECM), promoting tumor cell invasion, and downregulating antitumor immune responses via regulatory T-cell recruitment^7–9^. Via modulation of vascular endothelial growth factor (VEGF), transforming growth factor beta (TGFβ), and integrin α5β1 signaling, NRP-1 plays a role in multiple oncogenic path-ways in GB and is strongly associated with poor clinical outcomes^7–9^.

NRP-1 is known to bind several ligands relevant to GB pathophysiology. Among others, b1 domain of NRP-1 (b1-NRP-1) binds the C-terminal Arg residue of VEGF-A165 (Figure 1) ^9–13^. The C-terminal Arg residue of VEGF-A_165_ interacts with key residues of b1-NRP-1 (Tyr297, Asp320, Ser346, Thr349, and Tyr353)^7,14–15^. Mechanistically, the positively charged side chain guanidinium group of the C-terminal Arg residue forms a bidentate salt bridge with the conserved Asp320, while the C-terminal ⍺-carboxylate hydrogen-bonds to Ser346, Thr349, and Tyr353 residues^10^. The binding of the C-terminal Arg residue is further stabilized by cation-π and CH-π interactions^16–19^ with the aromatic side chains of Tyr297 and Tyr353 residues of b1-NRP-1^10,13^. Additionally, Thr316, and Trp411 residues of b1-NRP-1 are also crucial for high affinity binding of VEGF-A165^14^. On the basis of VEGF-A_165_ − b1-NRP-1 interactions, as inhibitors of the binding, several peptide analogs were designed^11–12,20^. Structural analysis of these peptides resulted in introduction of the C-end rule (CendR)^12^. The CendR is defined as a C-terminal peptide motif of R/KXXR/K sequence in which XX are any residues and the motif has a free ⍺-carboxylate end. The terminal basic residue, typically either Arg or Lys, docks deep into the conserved b1-NRP-1 binding pocket and competitively blocks the site^12^ The VEGF-A_165_ − b1-NRP-1 interaction enhances VEGF-A_165_ angiogenic signaling, making NRP-1 a key target for disrupting tumor vascularization therapeutically.

**Figure 1.**
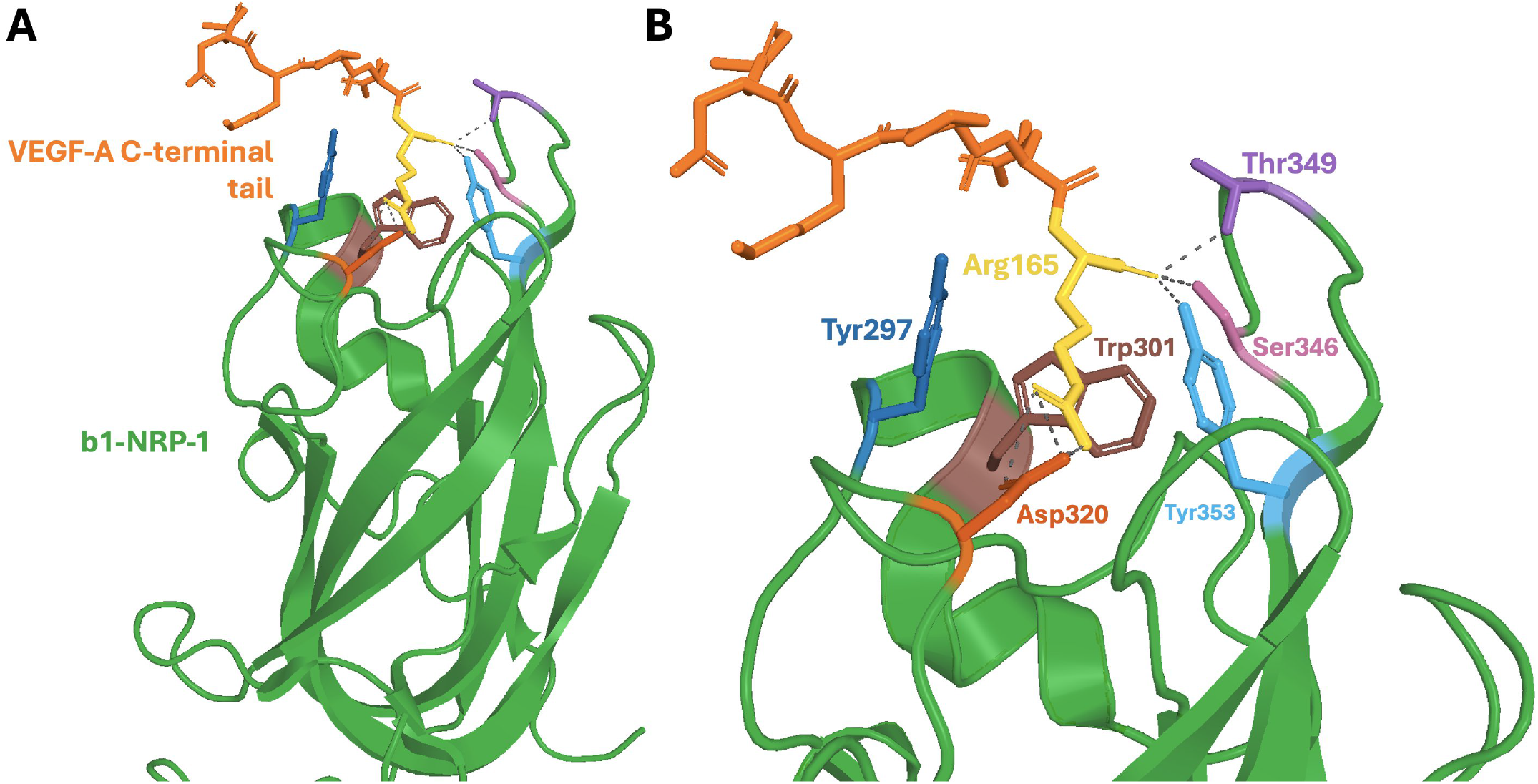
Structural basis of VEGF-A_165_ C-terminal (CendR) interaction with the b1-NRP-**1. A**, b1-NRP-1 domain (green) binds the C-terminal tail ofVEGF-A_165_ (orange), whose terminal Arg165 (yellow) inserts into the canonical CendR-binding pocket. **B**, zoomed-in view of the VEGF-A_165_ binding pocket. Six canonical pocket residues are shown as sticks and individually colored as Asp320 (vermilion), Tyr297 (blue), Tyr353 (sky blue), Ser346 (pink), Thr349 (purple), and Trp301 (brown). The VEGF-A_165_ C-terminal Arg165 binds the pocket through a salt bridge between its guanidinyl side chain group and Asp320, and through coordination of its free C-terminal carboxylate by Ser346 and Thr349 residues, and Tyr353; Tyr297, Tyr353, and Trp301 residues line the pocket. Grey dashed lines indicate polar contacts (≤ 3.6 Å). Structure was rendered from PDB 4DEQ^13^.

Laminin, a heterotrimeric ECM protein made up of α, β, and γ chains, is another key component of GB invasion pathophysiology. Laminins participate in cell adhesion and migration through integrin receptors, in glioma predominantly α3β1, and are often overexpressed, facilitating tumor cell migration and invasiveness^21–22^. NRP-1 regulates α5β1 integrin trafficking and adhesion. Laminins and NRP-1 both influence integrin-mediated adhesion and migration; it is, therefore, plausible that laminin-derived fragments might also interact with NRP-1. This possibility has an important therapeutic implication: peptide-based inhibitors from ECM proteins have the potential to disrupt tumor-promoting protein-protein interactions while also offering relatively high specificity and low toxicity^23–27^. Peptide modification such as cyclization, N-methylation, retroinverso modification and incorporation of non-canonical residues in peptides have greatly enhanced peptide protease resistance, stability, and blood-brain barrier (BBB) permeability, thereby expanding their applicability to CNS malignancies^24,27–28^.

Residues 929-933 fragment of the β1 chain of laminin, Tyr-Ile-Gly-Ser-Arg (peptide **1** in Table 1), was identified as a biologically active fragment^29–31^ inhibiting cell adhesion, migration, metastasis, and tumor cell attachment *in vitro*^29–31^. These landmark studies revealed that Tyr-Ile-Gly-Ser-Arg functions as a minimal recognition sequence for laminin-binding receptors and plays an important role in cellular interaction with the extracellular matrix. Structural analogs of peptide **1**, such as a retro analog with the reverse amino acid sequence (Ac-Arg-Ser-Gly-Ile-Tyr-NH_2_, peptide **2** in Table 1), and a retro-inverso analog in which the retro sequence (Ac-arg-ser-Gly-iletyr-NH_2_, peptide **3** in Table 1) (lower case letters designate D-amino acids), were developed^32^. These analogs aimed to strengthen proteolytic stability and enhance therapeutic potential^32–34^. Collectively, these works established the foundation of peptide **1** as an laminin-derived peptide fragment capable of modulating cancer cell adhesion and migration, laying the basis for peptide-based strategies targeting ECM-receptor interactions seen today.

**Table 1.** Primary structure of the peptides and AlphaFold 3 complex-prediction confidence scores.

| Peptide | Sequence | pTM | ipTM |
|---|---|---|---|
| **1** | Ac-Tyr-Ile-Gly-Ser-Arg-NH<sub>2</sub> | 0.95 | 0.84 |
| **2** | Ac-Arg-Ser-Gly-Ile-Tyr-NH<sub>2</sub> | 0.93 | 0.70 |
| **3**<sup>a</sup> | Ac-arg-ser-Gly-Ile-Tyr-NH<sub>2</sub> | - | - |

Although peptide **1** does not completely satisfy the R/KXXR/K CendR motif, it has a C-terminal Arg residue, a feature shared by canonical CendR ligands that bind to the b1-NRP-1^12^. The retro and retro-inverso peptides, as an anti-CendR analogs, contain an aromatic residue at their C-terminus. Our overall hypothesis is that peptide **1** and its analogs could either bind to the canonical VEGF-A165 binding site or associate with an allosteric site on the b1-NRP-1 domain. We also hypothesized that the C-terminal Tyr residue in the retro peptide **2** and retro-inverso pep-tide **3** as anti CendR analogs also can bind the b1-NRP-1 through π – π interactions^35–36^ with aromatic residues in the binding pocket. In this work, we aimed to elucidate whether peptide **1** and its analogs indeed bind to either the VEGF-A165 binding site or an allosteric site on the b1-NRP-1 and determine the structure, interactions and energetics of binding via *in silico* methods. We combined AlphaFold3.0 (AF3)^37^ structural prediction peptide − b1-NRP-1 complexes, all-atom molecular dynamics (MD) simulations, b1-NRP-1—peptide interaction analyses, and binding energy calculations by using FoldX^38–39^ to determine the potential of laminin-derived peptides as b1-NRP-1 binders. In addition to the computational analysis, measuring synthetic peptide binding to recombinant NRP-1 was performed using differential scanning fluorometry (DSF) assay. Presented results lay foundation for future studies to investigate the potential inhibition of NRP-1 activity using laminin-derived peptides.

## 3. Methods

### 3.1 Complex Prediction and Structure Preparation

AF3^37,40^ web server was used to generate three-dimensional (3D) structures of the peptide − NRP-1 complexes. Primary structure of the human b1-NRP-1 (residues 273–427) together with either peptide **1** or peptide **2** were submitted five times to the AF3 web server. For each prediction, five seeds were requested and the structure with the highest predicted template modeling (pTM) and interface predicted template modeling (ipTM) scores^41^ were saved for further studies. Using the YASARA^42–43^ software, in the retained complexes both the protein and peptides were N-terminally acetylated and C-terminally amidated to preserve the electronic structure of the backbone atoms. In addition, for the retro-inverso analogs the L-amino acids were replaced with their D-stereoisomer using the peptide **2** − b1-NRP-1 complex. Structures then were energy-minimized in the absence and then presence of explicit water using the AMBER-FB15 force field with Particle Mesh Ewald (PME) electrostatics under periodic boundary conditions^44–45^. Structures were saved in PDB format^46^ and used for subsequent MD simulations.

### 3.2 Molecular Dynamics (MD) Simulations

MD simulations were performed using GROMACS 2022.6 package^47^ and the CHARM-M36m force field^48^ as it was described in a previous publication^49^. Complexes were solvated with TIP3P water and placed in a dodecahedron box of a minimum solute-box edge distance of 1.2 nm. System charges were then neutralized to an ionic strength representative of physiological conditions of 150 mM NaCl^47^. His287 was modeled as the Nδ-H tautomer (HSD) and the Cys275-Cys424 disulfide bond was retained. Solvated complexes were submitted to 1000 steps of steepest-descent energy minimization, and then 1 ns constant number of particle, volume, and temperature (NVT) MD simulation at 300 K using the stochastic velocity rescale method^50^ with a temperature coupling constants (τ_T_) of 0.1 ps. Positions of backbone atoms were restrained with 1000 kJ mol nm force constant and integration timestep was 0.002 ps.

Equilibration using the stochastic velocity-rescale thermostat (τ_T_ = 0.1 ps) and the Berendsen barostat (τ_P_ = 2.0 ps, compressibility = 4.5 × 10**⁻⁵** bar**⁻**¹) under NPT ensemble at 310 K was accomplished with a 10 ns simulation time^50–51^. Production MD simulations of 1 µs were then executed in NPT ensemble using the stochastic cell-rescale (c-rescale) barostat (τ_P_ = 5.0 ps) and velocity-rescale thermostat (τ_T_ = 1.0 ps) at a temperature of 300 K^50,52^. Each simulation was performed with a 0.002 ps integration time step and LINCS constraints applied to all covalent hydrogen bonds^53^. Short-range van der Waals interactions were treated with a force-switch potential between 1.0 and 1.2 nm, while electrostatic interactions were computed using PME with a real-space cutoff of 1.2 nm^45^. Configuration, forces and energies were stored in intervals of 25 ps for subsequent analysis. A 5 μs of total production simulations per peptide and 15 μs of total production simulations were performed.

### 3.3 Trajectory Analysis

#### Equilibration assessment

For each peptide − b1-NRP-1 complex, the covariance matrix of the C_α_ atom movements was computed with the GROMACS *covar* utility. Equilibration was assessed in two ways. First, using the eigenvectors of the 100 largest eigenvalues, the configurational entropy was calcuted as a function of simulation time at a 2 ns sampling interval^54–57^. Second, the convergence of the sampled essential subspace was quantified as the normalized overlap between the covariance subspace of each growing trajectory segment and that of the full trajectory, evaluated at a 2 ns interval by the method of Hess^58^. The equilibrated portion of each trajectory used for subsequent analysis was determined from these phases.

#### Root mean square deviation (RMSD)

RMSD of C_α_ atom positions was computed for the b1-NRP-1 receptor, the respective pep-tide in each peptide-system, and the full complex to monitor structural stability during production simulation^47^.

#### Feature clustering

For each production trajectory, an in-house python script, using the MDAnalysis^59–60^ python library, was executed to perform feature clustering as follows. Inverse minimum inter-atomic distances between each C_α_ atom of b1-NRP-1 and each peptide C_α_ atom, as a feature, was calculated, so that close contacts dominate the representation. Dimensions of the resulting feature trajectories (155 × 5 = 775 features) were reduced by principal component analysis (PCA) of scikit-learn python library^61^. The first five principal components (PCs) trajectories were clustered by density-based hierarchical clustering (HDBSCAN^62^) method with a minimum cluster size of 5 % of frames and a minimum sample count of 1 %. Unassigned frames were labeled noise. The representative frame of each cluster was that nearest the cluster mean in PC subspace. Clusters were projected back onto the C_α_ RMSD of the original trajectories. Cluster representatives obtained by HDBSCAN were identified. Projected clusters onto the RMSD were used to identify the b1-NRP-1 pocket-associated windows and selected subsequent binding energy calculations using FoldX.

#### Canonical-pocket occupancy

Intervals during which the peptide occupied the VEGF-A_165_ binding pocket of b1-NRP-1, were identified from the clustering and interaction-frequency analyses. For each trajectory, occupancy of VEGF-A_165_ binding pocket of b1-NRP-1 was calculated as a ratio of designated cluster time-window and over all time of the equilibrated trajectory reported as mean ± SD across the five replicates per peptide. These same windows were used for FoldX binding energy calculations. This window-based metric quantifies how persistently the peptide resided in the pocket and is distinct from the FoldX interface frequencies below.

#### Binding frequencies and binding energies (ΔE_b_)

*AnalyseComplex* command of FoldX 5.1^38–39,63^ was used to sample at 1 ns intervals of the cluster time-window. For each sampled-frame FoldX reported interacting residues at the peptide – receptor interface^38–39^. For each receptor and peptide residue, using an in-house bash and python scripts, the interface frequency was the number of FoldX-evaluated frames in which the residue was at the interface divided by the total number of FoldX-evaluated frames in that trajectory, re-ported as mean ± SD across trajectories. For the receptor-wide interface profile, the terminal Ace- and -NH_2_ caps, were identified as a flexible-tail artifact and were excluded.

From the same sampled frames each structure was analyzed with the *RepairPDB* command of FoldX, and the binding energy (ΔE_b_) of the complex was computed with the *AnalyseComplex* command of FoldX^38–39^ using an in-house bash script. For each cluster-window, ΔE_b_ was averaged over all equilibrated frames; per-peptide values are reported as mean ± SD across pocket-associated windows. Since multiple windows can arise from the same trajectory, these window-level observations are not statistically independent and were interpreted descriptively.

#### Interaction analysis

Four classes of peptide – b1-NRP-1 interaction were quantified at 100 ps steps over each equilibrated interval of trajectories (Supplementary Table S1) using MDAnalysis. Only interchain contacts were considered. Reported occupancy is the percentage of equilibrated frames in which at least one interaction of that class was present, reported as mean ± SD across the five replicates.

#### Salt bridges

Ionic interactions between the Arg residue of the peptide and Asp320 of b1-NRP-1 were calculated directly from the production trajectories using VMD 2.0.0^64^ with an in-house Tcl script. For each frame after equilibration, distances between the Arg guanidinium nitrogen atoms (NE, NH1, NH2) and the Asp320 carboxylate oxygen atoms (OD1, OD2) were computed for each frame, applying minimum-image periodic boundary conditions. Ion-pair occupancy was the fraction of equilibrated frames with a minimum N···O distance ≤ 4.0 Å^65^. Bidentate geometry was evaluated separately and required NH1 and NH2 to contact different Asp320 carboxylate oxygens simultaneously, each within 3.5 Å. Values are mean ± SD across the five replicates per peptide.

#### Hydrogen bonds

Hydrogen bonds between the peptide and b1-NRP-1 were evaluated over the equilibrated portion of each trajectory. A hydrogen bond was counted when the donor-acceptor distance was ≤ 3.0 Å and the D—H···A angle was ≥ 160°. Only interchain donor-acceptor pairs were considered. Occupancy was the percentage of equilibrated frames containing at least one such interaction, reported as mean ± SD across the five trajectories.

#### Cation-π interactions

Cation-π interactions between the Arg of peptides and the aromatic residues (Tyr297, Trp301, Tyr353, Trp411) of b1-NRP-1 were evaluated over the equilibrated portion of each trajectory. For each frame, the distance from the Arg guanidinyl carbon (C_ζ_) to the aromatic ring centroid and the in-plane lateral offset of the cation were computed; a cation-π interaction was counted when the C_ζ_-to-centroid distance was ≤ 6.0 Å and the lateral offset was ≤ 2.3 Å^66–67^. For Trp residue, the six-membered benzene ring part of the indolyl ring was used. Occupancy was the percentage of equilibrated frames satisfying both criteria, reported as mean ± SD across the five trajectories.

#### CH-π interactions

CH-π interactions between the peptides and the b1 pocket aromatic residues were evaluated over the equilibrated portion of each trajectory. For each frame, the distance from a hydro-gen-bearing carbon to the aromatic ring centroid and the angle between the ring normal and the centroid-to-carbon vector were computed; a CH-π interaction was counted when the distance was ≤ 4.5 Å and the angle was ≤ 30°. For Trp, the six-membered ring of the indole was used. Occupancy was the percentage of equilibrated frames satisfying both criteria, reported as mean ± SD across the five trajectories.

### 3.4 Peptides and Recombinant NRP-1

Greater than 95% pure N- and C-terminally acetyl and amide protected, respectively, pep-tides were obtained from Peptide 2.0 Inc. (Chantilly, VA, USA). Extracellular domain of the recombinant human NRP–1 (UniProt: O14786) was obtained from ACROBiosystems, cat #NR1-H5228 (Newark, DE, USA), containing amino acid residues Phe22-Lys644.

#### NanoDifferential Scanning Fluorimetry (nanoDSF)

Assays were performed using the Prometheus Panta System (NanoTemper Technologies, München, Germany). Samples were prepared in 7.4 pH Tris buffer (20 mM Tris, 150 mM NaCl), with the final solution containing 1 µM protein and 1 – 100 µM concentration range of peptide (1, 3, 10, 20, 30, 45, 100 µM). Triplicate samples were loaded into Prometheus high sensitivity capillaries and exposed to a temperature gradient ranging from 15 °C to 85 °C, with a thermal ramping rate of 1.2 °C/min. Trp fluorescence was measured with 280 nm excitation and at a ratio of 350/330 nm emission with dual-UV detector. Assays were independently repeated three-times. Apparent binding affiinity (K_D,app_) was calculated using a local copy of the FoldAffinity software of the Online Data Analysis Platform for Molecular Biophysics web server^68–71^.

## 4. Results

### 4.1. AlphaFold3 complex prediction

Scores for AF3-predicted models of the peptide − b1-NRP-1 complexes listed in Table 1. pTM assesses confidence in the overall predicted fold (values > 0.5 indicate a model likely resembling the true structure), whereas ipTM assesses the accuracy of the modeled interface between subunits (> 0.8, confident; 0.6–0.8, intermediate; < 0.6, likely inaccurate). For peptide **1** the top-ranked model reached pTM of 0.95 and ipTM of 0.84, and for peptide **2** pTM of 0.93 and ipTM of 0.70; The scores were reproduced across all five independent AF3 simulations per peptide. The high pTM values indicate confident overall folds for all complexes. At the interface, peptide **1** (ipTM = 0.84) was predicted with high confidence, whereas peptide **2** (ipTM = 0.70) fell in the intermediate range, suggesting peptide **2** had a plausible but less certain placement of the retro analog in the pocket. These predictions provided reasonable structures for subsequent MD simulations, with the retro structure of peptide **2** (and, by inheritance, the retro-inverso peptide **3**) analog interface predicted less confidently than that of peptide **1**. Peptide **3** was generated from a peptide **2** by replacing all L-amino acids with their D-enantiomer and therefore inherits the peptide **2** confidence estimates rather than having an independent AF3 score.

### 4.2. Trajectory stability of the peptide—b1-NRP-1 complexes

Visual inspection of the trajectory-specific configurational-entropy plots (Supplementary Figure S1) identified sustained lower-entropy plateaus during the initial portion of each production simulation. The selected equilibrated cutoffs occurred at a mean and ± standard deviation (SD) of 44.6 ± 18.9 ns for peptide **1**, 46.1 ± 30.9 ns for peptide **2**, and 33.5 ± 5.9 ns for peptide **3** (Supplementary Table S1; Supplementary Figure S1). RMSD and all other subsequent analyses were, therefore, performed for each trajectory through the equilibration phase of the 1000 ns production simulation. Following equilibration, the b1-NRP-1 domain remained structurally stable, with mean ± SD C_α_ RMSD values of 0.227 ± 0.025 nm for peptide **1**, 0.213 ± 0.009 nm for peptide **2**, and 0.247 ± 0.060 nm for peptide **3**. Peptide RMSD values were higher, consistent with the expected flexibility of short and partially solvent-exposed pentapeptides, with equilibrated mean and SD values of 0.386 ± 0.040 nm for peptide **1**, 0.397 ± 0.026 nm for peptide **2**, and 0.378 ± 0.056 nm for peptide **3** (Figure 2). These results indicate that the b1-NRP-1 domain remained stable across all complex systems and that differences in pocket occupancy were not driven by b1-NRP-1 instability.

**Figure 2.**
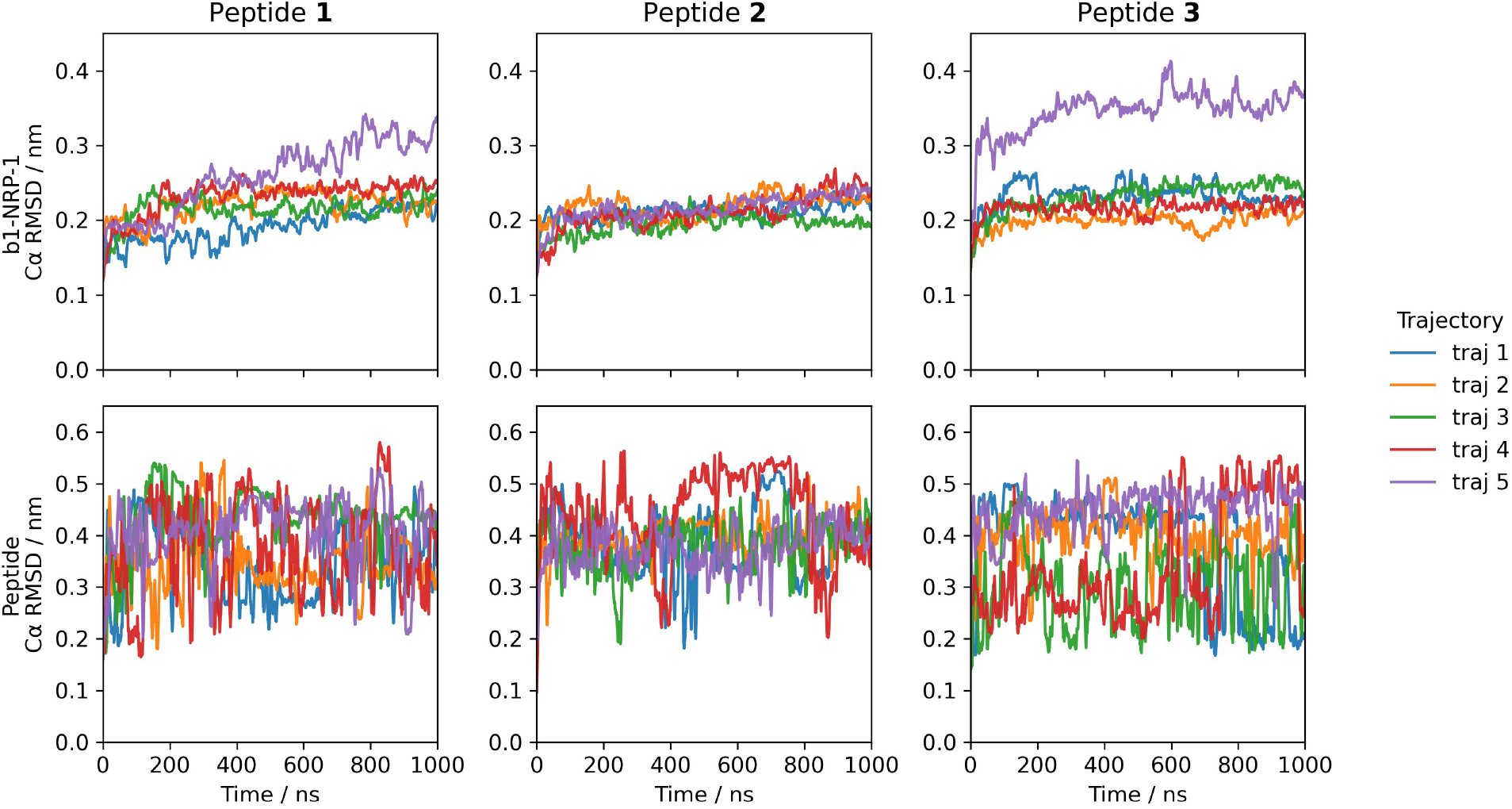
Structural stability of the peptide – b1-NRP-1. C_α_ RMSD of the b1-NRP-1 (top row) and peptide (bottom row) across five independent trajectories for each system. The b1-NRP-1 remained broadly stable across the simulations, with equilibrated mean RMSD values of approximately 0.21–0.25 nm, supporting the interpretation that differences in pocket occupancy were not attributable to gross b1-NRP-1 instability. A potential outlier of trajectory 5 for b1-NRP-1 in the peptide system was observed. Peptide RMSD values were higher and varied more, consistent with the expected flexibility of solvent-exposed regions of the pentapeptides.

### 4.3. Feature clustering

For each trajectory the leading PCs captured the majority of the inverse contact variance, and clustering resolved a small number of discrete conformational states together with unassigned (noise) frames. Projecting cluster membership onto the complex C_α_ RMSD trace separated a compact, low RMSD pocket-bound state from higher RMSD states in which the peptide dissociated (Figure 3). In representative trajectories, peptide **1** settled into a single densely populated pocket-bound cluster, whereas peptide **2** sampled several higher RMSD states without a dominant pocket-bound cluster and peptide **3** occupied a stable pocket-bound state early before dissociating, which is consistent with the occupancy differences quantified below (Figure 4). These pocket-associated clusters defined the windows used for the occupancy and FoldX analyses, providing a trajectory specific basis for identifying pocket-bound intervals.

**Figure 3.**
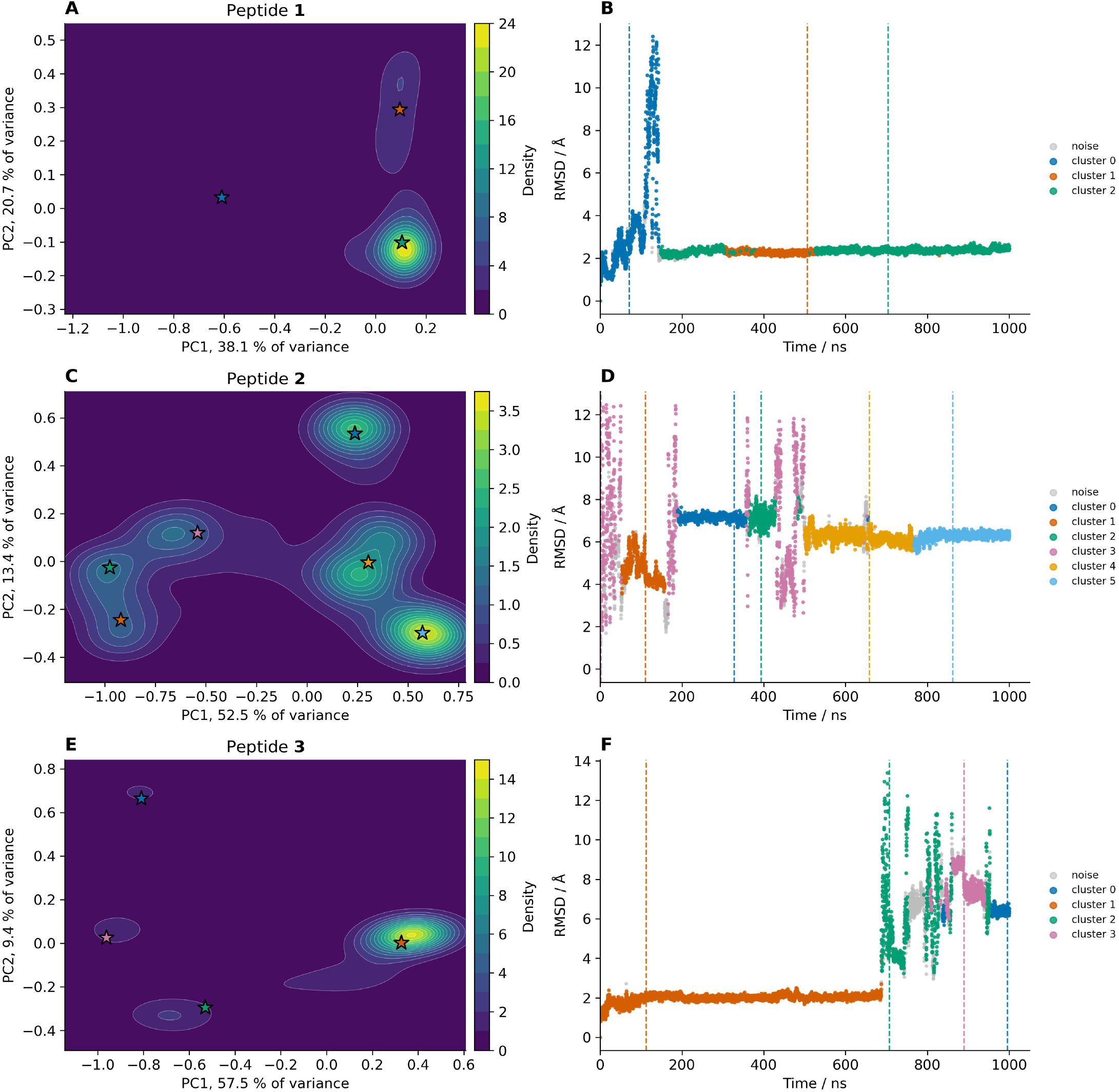
Inverse contact-based conformational feature clustering for a representative trajectory of each peptide. Panels are for peptide **1**, trajectory 3; peptide **2**, trajectory 1; peptide **3**, trajectory 1. **A**, **C**, **E**, kernel density surface of the frames in the first two principal components of the peptide – b1-NRP-1 contact space (percent variance on axes) for peptides **1**, **2**, and **3**, respectively; stars mark the HDBSCAN cluster representatives. **B**, **D**, **F**, C_α_ RMSD of the complex relative to the first frame, colored by HDB-SCAN cluster membership (gray, clustering noise); vertical dashed lines mark the time frame of the cluster representatives. This RMSD is computed over all complex Cα atoms and is therefore dominated by peptide displacement once the receptor is equili-brated (compare to. the receptor only RMSD in Figure 2); large deviations correspond to the peptide leaving the pocket. Peptide **1** (A, B) settles into a single densely populated, low RMSD pocket-bound state; peptide **2** (C, D) samples multiple higher RMSD states without a dominant pocket-bound cluster; peptide **3** (E, F) occupies a stable low RMSD state early before dissociating. These clusters defined the pocket-associated windows used for the occupancy (Figure 4) and FoldX analyses.

**Figure 4.**
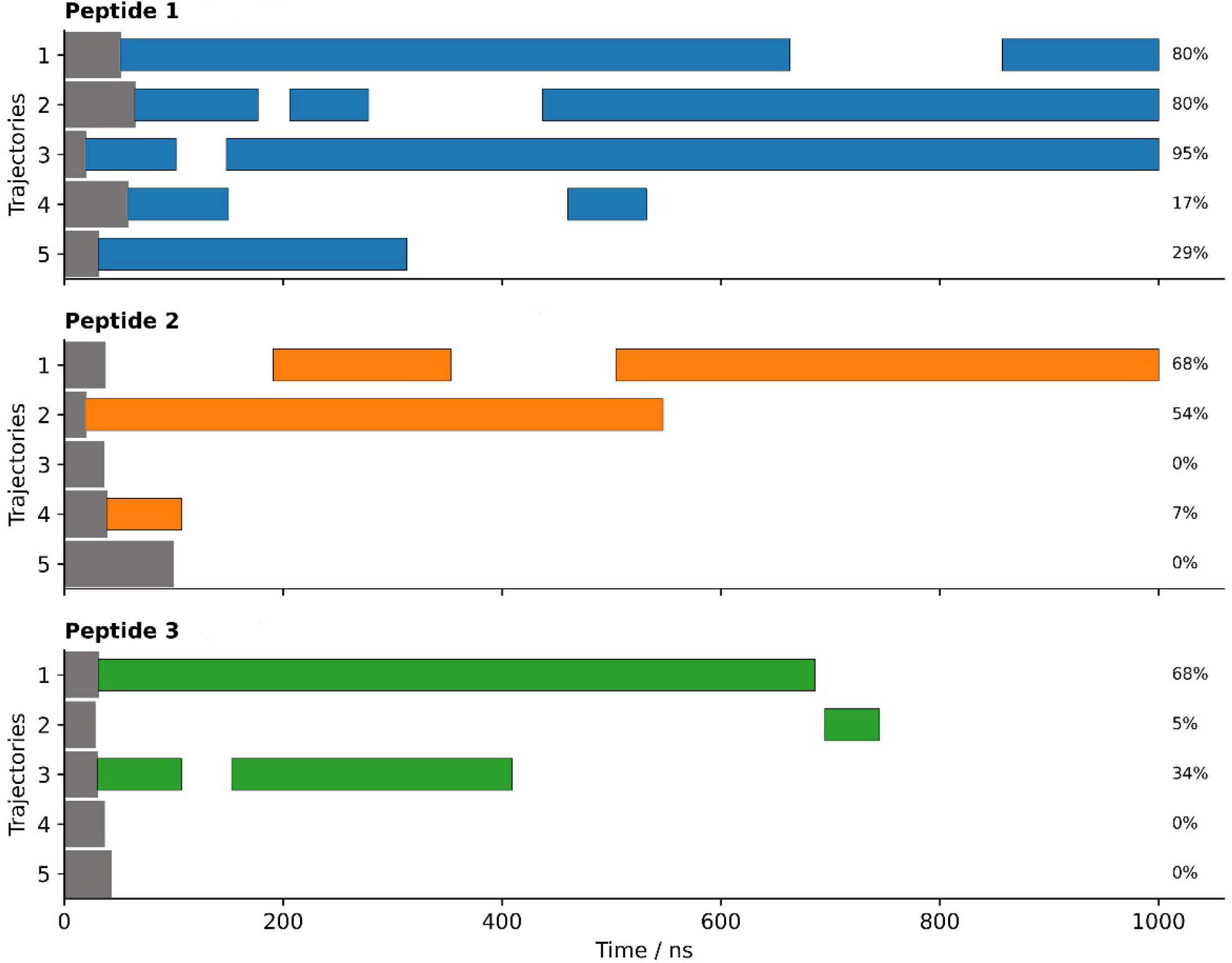
Residence of each peptide in the canonical VEGF-A_165_ binding pocket across five independent trajectories, over the equilibrated interval. Dark grey bars indicate the trajectory-specific pre-equilibration intervals (Supplementary Table S1), which were excluded from analysis. Blue, orange and green bars mark the equilibrated intervals during which peptide 1, 2 and 3, respectively, was bound within the pocket; Right side indicates the percentage of b1-NRP-1 binding pocket occupancy in the trajectory (summed equilibrated pocket-bound duration divided by the over all equilibrated duration). The mean ± SD occupancy across five trajectories are 60.3 ± 34.6%, peptide 1; 25.9 ± 32.7%, peptide 2 and 21.4 ± 29.5%, peptide 3.

Of the three peptides, peptide **1** demonstrated the highest pocket occupancy at 60.3 ± 34.6% (Figure 4). The retro analog peptide **2** and retro-inverso analog peptide **3** showed lower pocket occupancies of 25.9 ± 32.7% and 21.4 ± 29.5%, respectively (Figure 4). Peptide **1**, therefore, maintained more persistent occupancy of the canonical VEGF-A_165_-binding pocket than either analog, although substantial between trajectory variability was observed.

Across the full b1-NRP-1 domain, the residues most frequent interactions were concentrated in the canonical VEGF-A_165_ pocket (shaded green), which stood out above the surrounding receptor surface for all three peptides (Figure 5A).

**Figure 5.**
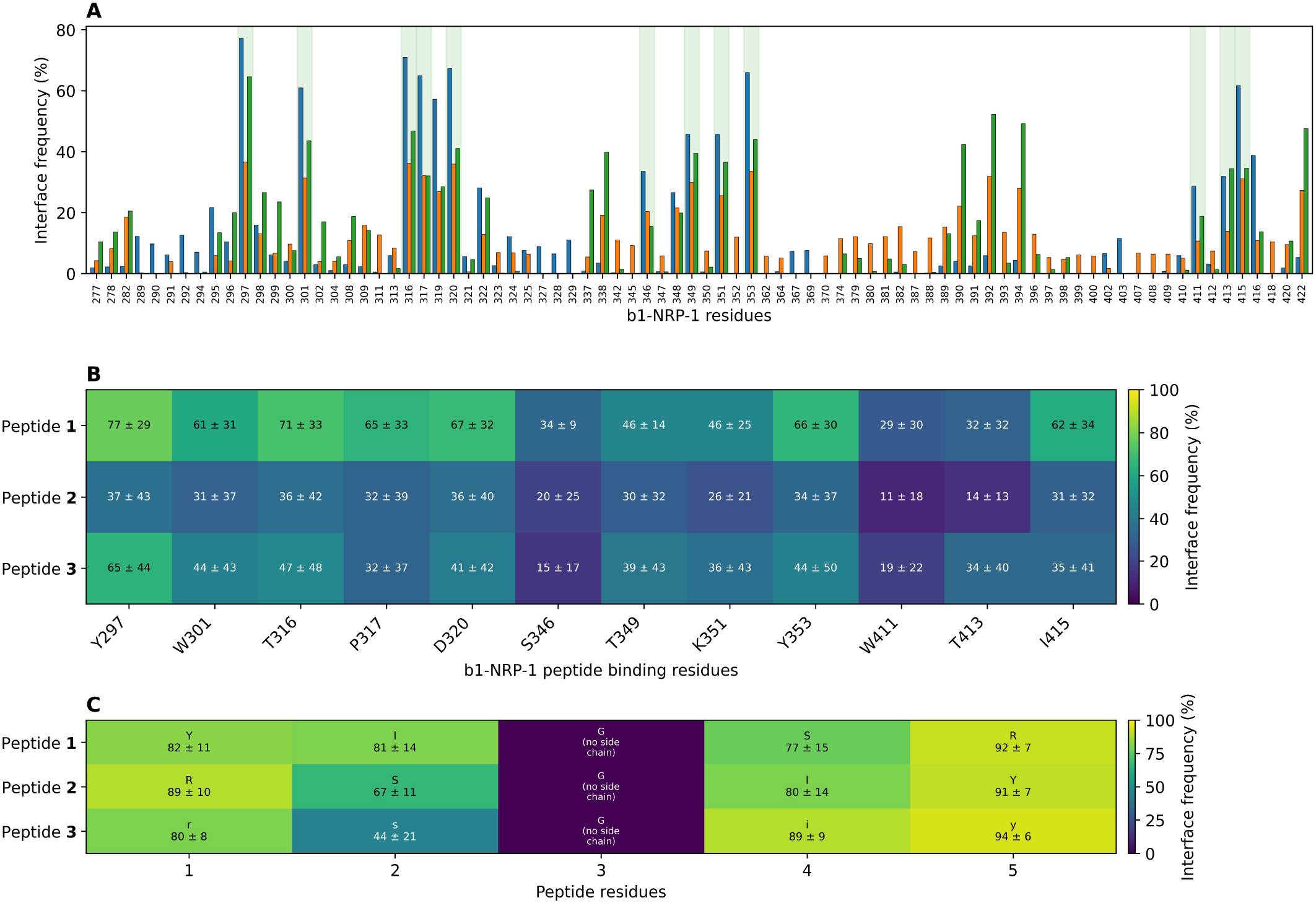
Interface-residue frequencies across the b1-NRP-1 domain determined by FoldX. **A**, binding frequencies of b1-NRP-1 residues determined by *AnalyseComplex* command for peptide **1,** blue; peptide **2**, orange; peptide **3**. green. Interface frequency was counted for b1-NRP-1 residues reaching ≥ 5% in at least one peptide. Terminal caps and residues 273–276 are excluded. The VEGF-A_165_ binding residues are shaded green. Interface frequency is the fraction of FoldX-evaluated frames in which a residue was at the interface, computed per trajectory and reported as mean ± SD across trajectories. This interface frequency is conditional on the peptide being in a FoldX-evaluated pocket state and therefore describes which residues form the interface when the peptide is interacted with. **B,** Binding interface frequency of the twelve canonical pocket residues of b1-NRP-1 with peptides. **C**, Binding interface frequency of peptide residues. The central Gly residue has no side chain and is not identified by FoldX as interacting residue. Across FoldX-evaluated pocket conformers, the canonical pocket residues form the interface most frequently for peptide **1**, while terminal and aromatic tyrosyl residues of each peptide also interacted at high frequency.

For peptide **1**, the amino acid residues most frequently at the interface were the C-terminal Arg5 (92 ± 7% of FoldX-evaluated frames), Tyr1 (82 ± 11%), Ile2 (81 ± 14%), and Ser4 (77 ± 15%) (Figure 5C). On the receptor side, peptide **1** most frequently interacted with Tyr297 (77 ± 29%), Thr316 (71 ± 33%), Asp320 (67 ± 32%), Tyr353 (66 ± 30%), and Pro317 (65 ± 33%), residues within or adjacent to the canonical VEGF-A_165_-binding pocket of b1-NRP-1 (Figure 5B)^11–12,20^_._

When in a pocket-associated state, the peptides **2** and **3** interacted with the same interface residues at broadly comparable frequencies, indicating that the peptides differ mainly in how of-ten they occupy the pocket (Figure 4) rather than in how they interact with b1-NRP-1 residues. For peptide **2**, the most frequent peptide interface residues were Tyr5 (91 ± 7%), Arg1 (89 ± 10%), and Ile4 (80 ± 14%); for peptide **3**, D-Tyr5 (94 ± 6%), D-Ile4 (89 ± 9%), and D-Arg1 (80 ± 8%) (Figure 5C). On the b1-NRP-1 side, interface frequencies for the analogs were more variable across trajectories (for example, Tyr297 37 ± 43% for peptide **2** and 65 ± 44% for peptide **3**), reflecting their smaller number of pocket-associated windows and substantial between-trajectory variability (Figure 5B). Although peptide **1** occupied the canonical pocket more persistently than either analog (Figure 4), the analogs interacted with a similar set of interface residues when bound. Representative pocket-bound conformations of the three peptides illustrate this shared interactions (Figure 6).

**Figure 6.**
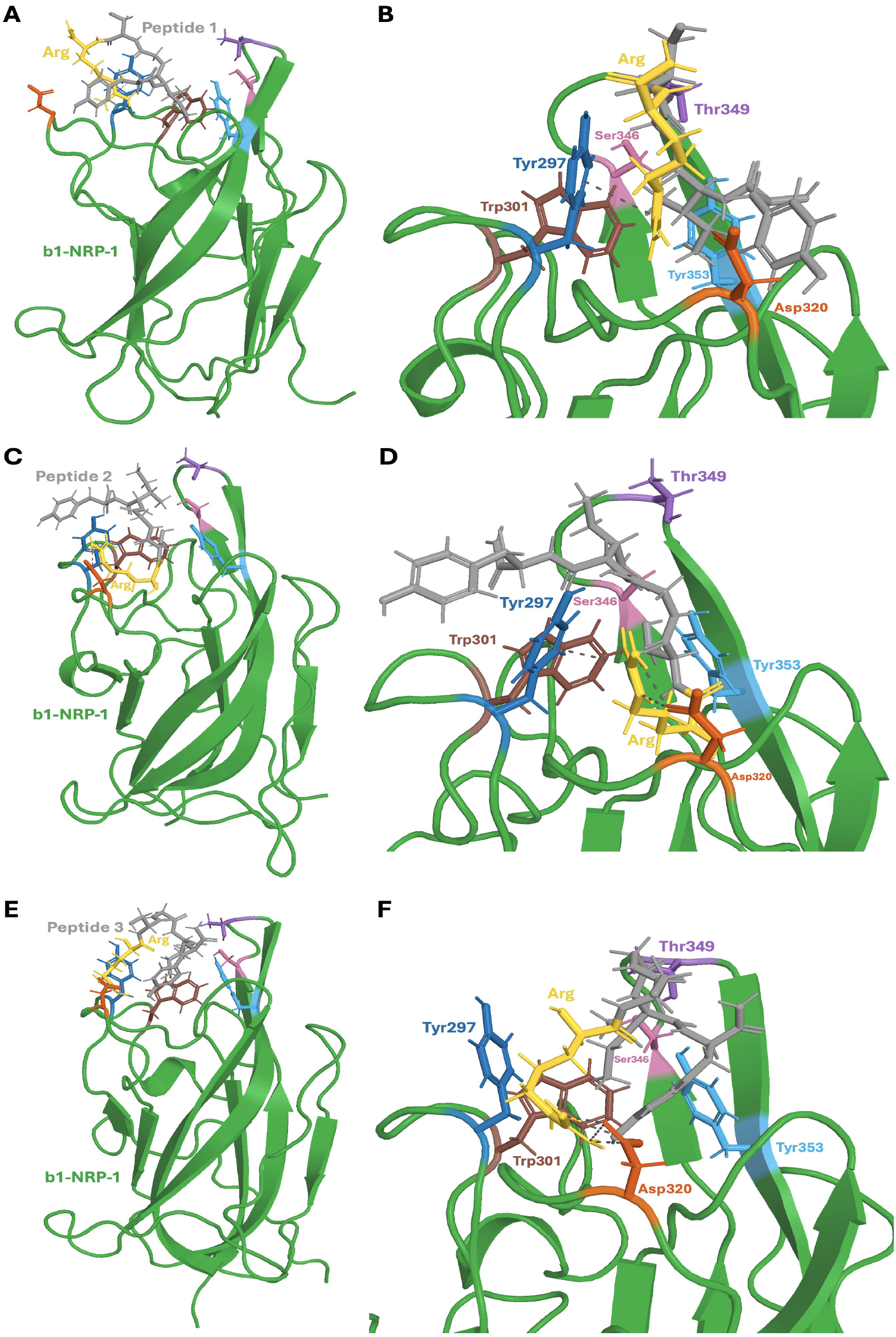
Lowest binding energy (ΔE_b_) structures of the three peptides in the b1-NRP-1 pocket. Peptide 1, A and B; peptide **2**, **C** and **D**; peptide **3**, **E** and **F**. Each structure are aligned to a common view of the b1-NRP-1 domain (green); panels **A**, **C**, and **E** show the overall domain. In panels **B**, **D**, and **F** structures are zoomed in to the binding interface. Each peptide is shown in gray with its Arg shown in yellow (Arg5 for peptide **1**; Arg1 for the peptide **2** and **3**). For the six canonical pocket residues the same colors are used as in Figure 1: Asp320 (vermillion), Tyr297 (blue), Tyr353 (sky blue), Ser346 (pink), Thr349 (purple), and Trp301 (brown). Grey dashed lines denote side-chain interactions: the Arg guanidinium–Asp320 salt bridge (side-chain N···O ≤ 4.0 Å) and Arg–Tyr cation-π-compatible geometry (guanidinium Cζ to aromatic-ring centroid ≤ 6.0 Å and ≤ 2.3 Å lateral offset). In these representative frames, the Arg residues of peptide **1** (B) and peptide **2** (D) sample cation-π-compatible geometry with Tyr297, whereas peptide **3** (F) instead shows the Arg–Asp320 salt bridge; no single interaction is present in every frame, consistent with the transient, interchangeable interactions described in the text. Because all peptides were simulated C-terminally amidated form, they lack the free α-carboxylate of canonical CendR ligands, so interactyions are mediated by the Arg side chain and aromatic-pocket contacts rather than the canonical carboxylate network shown for VEGF-A_165_ in Figure 1. When bound, the three peptides interact with a similar set of pocket residues, consistent with the interface analysis (Figure 5); they differ chiefly in how often they occupy the pocket (Figure 4).

### 4.4 FoldX binding energies (ΔE_b_)

ΔE_b_ calculations by FoldX binding energy estimates were done for 10, 4, and 4 pocket-as-sociated structural windows for peptide **1**, peptide **2**, and peptide **3**. For each window, ΔE_b_ was averaged over all equilibrated frames at 1 ns sampling. Because multiple windows could originate from the same trajectory, these observations were not statistically independent, therefore, they were interpreted descriptively, and no statistical comparison was carried out. The mean estimated ΔE_b_ and SD across windows were −3.42 ± 1.52 kcal mol^-1^ for peptide **1** (n = 10), −4.11 ± 1.07 kcal mol^-1^ for peptide **2** (n = 4), and −3.62 ± 2.26 kcal mol^-1^ for peptide **3** (n = 4) (Figure 7). Although the three peptides differed in pocket occupancy, their FoldX ΔE_b_ values were similar in magnitude and overlapped substantially.

**Figure 7.**
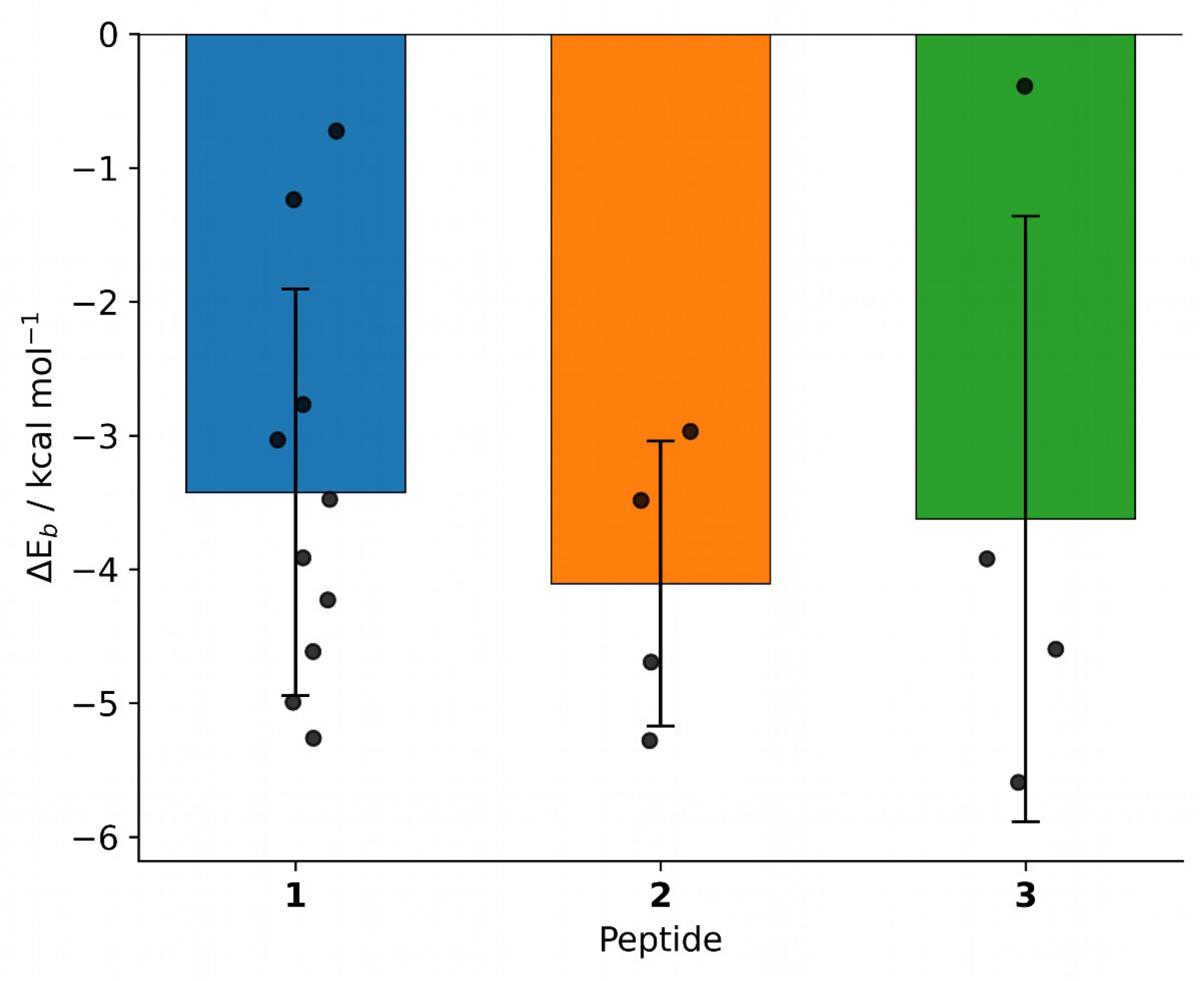
FoldX binding-energy estimates (ΔE_b_) for the peptide – NRP-1 b1 complexes. For each peptide, ΔE_b_ was computed with FoldX *AnalyseComplex* command after executing FoldX *RepairPDB* command; bars show the mean ± SD (Peptide **1** n = 10, peptide **2** n = 4, peptide **3** n = 4 windows), with individual window means indicated as points. The three peptides have ΔE_b_ estimates of similar magnitude that overlapped substantially.

### 4.5. Residue interactions in the b1-NRP-1 − peptide complexes

Peptide **1** interacted with the Asp320 residue of the VEGF-A_165_ binding pocket most persistently, including a canonical bidentate salt bridge, requiring both terminal guanidinyl NH groups to contact different Asp320 carboxylate oxygens simultaneously, and it was populated in 19.3 ± 20.1% of frames for peptide **1**, 10.7 ± 23.3% for peptide **2**, and 0.4 ± 0.7% for peptide 3. Under the standard ion-pair criterion the Arg-Asp320 contact occupied 32.7 ± 20.9%, 17.4 ± 22.7% and 7.2 ± 10.7% of frames respectively. Considering all acidic residues of b1-NRP-1, at least one ionic contact was present in 40.0 ± 17.0%, 36.2 ± 15.7% and 22.4 ± 6.4% of frames (Supplementary Figure S3A), with Asp320 the most frequent acidic partner in every system. Mean Arg-Asp320 occupancies followed the same rank order as canonical-pocket occupancy, although substantial between-trajectory variability was present.

At least one interchain hydrogen bond was present in 57.0 ± 11.4% of frames for peptide **1**, 55.9 ± 8.0% for peptide **2** and 43.2 ± 7.9% for peptide **3** (Supplementary Figure S3B). As with the ionic contacts, the dominant partner was Asp320: the Arg-Asp320 hydrogen bond occupied 20.0 ± 17.5% of frames for peptide **1**, 12.0 ± 17.0% for peptide **2** and 5.2 ± 7.2% for peptide 3. The peptidyl Arg residue also hydrogen-bonded to the Pro317 and Glu319 backbone at lower frequency (≤ 5.5% in every system). Hydrogen bonding was, therefore, more frequently populated than the ionic interaction in all three systems.

Since the VEGF-A_165_ binding pocket is also lined by aromatic residues, we examined whether the Arg residue of the peptides participated in cation-π interactions with them. For peptide **1**, the guanidinyl side chain group of Arg residue interacted with both Tyr297 (5.9 ± 6.5% of equilibrated frames) and Tyr353 (2.6 ± 2.3%); arrangements with Trp301 and Trp411 were negligible (≤ 0.4%). Cation-π interaction of Arg residue with Try297 was sampled at a similar mean frequency by peptide **2** (5.0 ± 6.8%), whereas, cation-π interaction with Tyr353 was more frequently sampled by peptide **1** than by either analog (peptide **2** 0.3 ± 0.5%; peptide **3** 0.3 ± 0.6%); peptide **3** showed little cation-π compatible sampling at either residue (Tyr297 0.7 ± 0.9%). As with the Asp320 salt bridge, these interactoins were transient rather than persistently maintained, and the large between-trajectory variability (SD comparable to or exceeding the mean, n = 5) precludes ranking the peptides by any single aromatic contact (Supplementary Figure S2, Supplementary Figure S3C).

CH-π interactions were the most frequently populated of the four described interaction classes, occupying 66.4 ± 20.4% of equilibrated frames for peptide **1**, 57.1 ± 10.7% for peptide **2** and 52.2 ± 13.1% for peptide **3** (Supplementary Figure S3D). For peptides **1** and **2**, Arg-Tyr297 CH-π contacts occurred in 30.5 ± 25.1% and 12.0 ± 20.0% of equilibrated frames respectively, with Cβ and Cγ methylene groups providing the predominant contacts (peptide **1**: Cβ 22.2 ± 22.6%, Cγ 23.2 ± 19.6%, Cδ 7.7 ± 4.1%; peptide **2**: Cβ 7.3 ± 12.7%, Cγ 8.4 ± 16.3%, Cδ 5.5 ± 8.9%; a frame may satisfy the criterion for more than one carbon, so these do not sum to the total); Tyr297 is the same residue that supported their cation-π interactions. In peptide **3** the Arg-Tyr297 contact was largely absent and the aromatic contacts were found with Ile4 interacting with Trp301 (11.1 ± 19.8%) and Phe335 (7.0 ± 9.5%). Taken together, the interaction analyses repeatedly implicated Asp320 and Tyr297 as major components of canonical-pocket interactions, while peptide **3** showed a shifted aromatic-contact pattern. Retro-inverso modification did not abolish interactions of the b1-NRP-1 pocket residues with peptide **3**, but redistributed its contacts away from the Arg-centred pattern of peptides **1** and **2**.

### 4.6. nanoDSF

nanoDSF thermal-shift assays yielded apparent dissociation constants (K_D,app_) of 20.52 ± 12.63 µM for peptide **1**, 12.72 ± 4.63 µM for peptide **2**, and 33.32 µM (Table 2). Peptide **2** had the lowest K_D,app_, the highest apparent affinity of the three peptides, in agreement with the FoldX calculations, in which peptide **2** also gave the most favorable mean ΔE_b_ (-4.11 ± 1.07 kcal mol^-1^; Figure 7). Peptides **1** and **3** differed in mean FoldX ΔE_b_ by only ∼0.2 kcal mol^-1^ (-3.42 ± 1.52 and -3.62 ± 2.26 kcal mol^-1^, respectively), well within the corresponding standard deviations, so the FoldX ΔE_b_ estimates did not differentiate these two peptides. Whereas in nanoDSF assay, peptide **1** had higher affinity apparent affinity than peptide **3**. The two independent approaches, therefore, agree on the most favorable peptide and are not in conflict for the remaining pair, which falls within the resolution of the FoldX ΔE_b_ estimates. This qualitative concordance between an independent biophysical measurement and the energetic scoring increases supports the computational results.

**Table 2.** Apparent binding affinities (K_D,app_) of peptides. Values are listed as mean ± SD

| Peptide | K<sub>D,app</sub> ± SD / µM |
|---|---|
| **1** | 20.52 ± 12.63 |
| **2** | 12.72 ± 4.63 |
| **3** | 33.32<sup>a</sup> |

<sup>a</sup>Two experiments with triplicate measurements of each.

## Discussion

In the present study we examined whether the laminin fragment peptide **1** and its structural analogs, peptide **2** and peptide **3**, interact with the b1-NRP-1 domain at or near the canonical VEGF-A_165_ binding pocket or an alternative allosteric site^11–12,20^. From AF3-based complex prediction, long-timescale molecular dynamics simulation, and residue-level contact analysis, pep-tide **1** showed the highest pocket occupancy within the VEGF-A_165_ binding pocket^37,47^. All three laminin-derived peptides contacted residues within the b1-NRP-1 pocket; peptide **1** demonstrated substantially greater pocket occupancy than either the retro or retro-inverso analog. No definite allosteric site was identified. The only substantial off-pocket interactions involved region at bothe the N- and C-termini of the b1-NRP-1 domain, which is an artifact, since these residues at either side are part the larger ectodomain og NRP-1. Present findings support our hypothesis that peptide **1** can associate with the VEGF-A_165_ binding region of b1-NRP-1 and suggest that laminin-derived fragments may represent a useful starting point for ECM-inspired NRP-1 antagonist development^7,20,23–24,29–30^_._

Although substantial between-trajectory variability was observed for all three peptides, peptide **1** had higher mean VEGF-A_165_ binding pocket occupancy compared with peptide **2** and peptide **3**. Peptide **1** occupied the pocket for 60.3 ± 34.6% of the equilibrated simulation, whereas peptide **2** and peptide **3** occupied it for only 25.9% ± 32.7% and 21.4% ± 29.5%, respectively. This is consistent with persistent association of peptide **1** with the canonical b1-NRP-1 pocket rather than nonspecific surface contact. Since the b1-NRP-1 domain mediates binding to VEGF-A_165_ and other described CendR-like peptides, the persistent association with this region is mechanistically relevant for potential inhibition^11–12,20^. Prior studies have shown that the VEGF-A_165_ binding site can be blocked using peptide inhibitors that follow the CendR rule, supporting the broader potential of peptide-based antagonism of NRP-1^12,20^. Therefore, the increased persistence of peptide **1** within the pocket supports the investigation of peptide **1** as a candidate scaffold for optimization.

Feature clusterings provided a trajectory-specific criterion for identifying pocket-bound trajectory intervals, and the resulting state maps reinforced the occupancy analysis: peptide **1** populated a single, stable pocket-bound state, whereas the peptides **2** and **3** either failed to settle into a dominant pocket-bound cluster or dissociated after an initial bound period. This convergence of an unsupervised clustering result with the window-based occupancy metric supports the conclusion that the peptides differ primarily in how persistently they hold a common binding mode, with peptide **3** additionally showing an altered aromatic-contact pattern likely due to its retro-inverso modification.

The higher occupancy persistence of peptide **1** is most likely due to its partial similarity to CendR-type binding behavior. CendR rule describes peptides with a C-terminal R/KXXR/K motif where the terminal basic residue, usually Arg or Lys, interacts with the b1-NRP-1 binding pocket^11–12^. Peptide **1** does not completely satisfy the canonical CendR motif, since it has a C-terminal amide Arg residue, therefore, did not possess the free α-carboxylate required for canonical CendR recognition. Similarities to the VEGF-A_165_/CendR binding mode were accordingly interpreted in terms of Arg side-chain localization and aromatic-pocket engagement rather than full reproduction of the canonical CendR interaction network. In contrast, the retro and retro-in-verso analogs, peptide **2** and peptide **3** respectively, have an aromatic residue at the C-terminus^32^. Arg5 residue of peptide **1** of maintained frequent contact with b1-NRP-1 pocket residues, including Asp320 and nearby aromatic residues of Tyr297 and Tyr353, which are part of the canonical VEGF-A_165_ – b1-NRP-1 pocket interactions^11,20^. While the terminal Arg residue of peptide **1** may contribute most to stable pocket retention when bound, Tyr residue of the analogs **2** and **3** also contribute to binding (Figure 6). However, they occupy the canonical pocket less often (Figure 4). This implies that the sequence and terminal positioning of residues may be important for sustained interactions of the b1-NRP-1 pocket with peptide **1** and its analogs^32–34^.

When peptide **1** occupied the canonical pocket, its terminal Arg residue displayed associations seen in canonical CendR recognition: interactions with Asp320, frequent contact with Tyr297, and transient cation-π interactions with Tyr297 and Tyr353 (Figure 6; Supplementary Figure S2). The Asp320 salt bridge was intermittent, peptide **1** maintained a canonical bidentate arrangement in roughly one fifth of equilibrated frames, and the cation-π interactions were transient, yet peptide **1** showed greater overall canonical-pocket residence than either analog. This is consistent with a dynamic interaction network at the pocket rather than a single persistent noncovalent bond. Across the interaction analyses, engagement of the Asp320/Tyr297-centred pocket was most persistent for peptide **1** and least persistent for peptide **3**, while the retro-inverso analog also exhibited a shifted aromatic-contact pattern.

FoldX ΔE_b_ estimates failed to clearly distinguish peptide **1** from its analogs. The mean ΔE_b_ values were similar in magnitude for each of the three peptides, with peptide **2** and peptide **3** having slightly more negative average values despite the lower pocket occupancy of the analogs previously seen. Therefore, the static nature of ΔE_b_ estimates alone do not fully describe peptide behavior in the b1-NRP-1 pocket. FoldX is an empirical scoring method that encompasses common energetic forces, including van der Waals interactions, hydrogen bonding, electrostatics, solvation, entropy, and steric penalties, but it does not fully capture more subtle interactions such as cation–π interactions, anion–π interactions, water-mediated contacts, or long-range electrostatic effects in an explicitly solvated environment^38–39,66,72–75^. This limitation may be particularly relevant for peptide **1**, where the C-terminal Arg residue interacts with both acidic and aromatic residues in the b1-NRP-1pocket^11–12,20^. For small, flexible peptides such as the peptides studied here, interactions with shallow, solvent-exposed binding surfaces may be better described via dynamic persistence and contact frequency than energy estimates from representative structures^47^. Therefore, the incongruity between FoldX energies and pocket occupancy is not evidence against peptide **1** binding, but rather it serves as an indication that dynamic trajectory-based metrics are necessary for evaluating peptide-protein interactions and that further energetic based studies using different methods *in silico* and *in vitro* should be performed.

The nanoDSF measurements provide experimental support for interaction between the peptides and recombinant NRP-1, extending the computational observations beyond an exclusively *in silico* preditiction. Peptide **2** showed the lowest K_D,app_ (12.72 ± 4.63 µM) and also had the most favorable mean ΔE_b_, giving qualitative support between the experimental assays and energetic calculations. The rank order of K_D,app_, however, was different from that of the ΔE_b_. Pep-tide **2** ranked above peptide **1** above peptide **3** by K_D,app_, whereas, ΔE_b_ separated peptides **1** and **3** by only ∼0.2 kcal mol^-1^, well within their standard deviations, and therefore did not differentiate them.

The decreased persistence of peptide **2** and peptide **3** binding is notable as retro and retroinverso peptide design is commonly used to improve proteolytic stability while maintaining biological activity^32–33^. However, in this study, reversing the sequence and applying retro-inverso stereochemistry reduced the percent pocket occupancy of the peptides **1** and **2** into the VEGF-A_165_ binding pocket. This does not exclusively mean that these analogs would lack biological activity, but it implies that direct mimicry of peptide **1** binding to NRP-1 is potentially sensitive to residue order, chirality, and terminal residue placement. Retro-inverso analogs can preserve specific global side-chain topology of a parent peptide, but they do not necessarily reproduce the same three-dimensional interaction pattern^33–34^. For peptide bindings to b1-NRP-1, C-terminal basic residue, either as free acid or amidated, may be more important for strong binding than having aromatic residue at that position^11–12,20^.

The present finding are in concordance with the biological background of peptide **1** as a laminin-derived fragment that participates in extracellular matrix interactions. Peptide **1** was first identified as a short laminin β1-chain sequence with the ability to modulate cell adhesion, migration, tumor cell attachment, and metastasis^29–31^. Since GB invasion is mediated via extracellular matrix remodeling, integrin signaling, angiogenesis, and cell-surface co-receptor activity, an interaction between a laminin-derived peptide and NRP-1 is biologically plausible^1,3,7–9,21–22^. NRP-1 potentiates VEGF signaling and α_5_β_1_ integrin-related pathways, while laminins and their derived peptides are involved in cell adhesion and migration via integrin-associated mechanisms as well ^8–9,21,29–30^. Therefore, the findings suggest peptide **1** binding to the b1-NRP-1 pocket represents a possible molecular connection between ECM-derived peptide motifs and NRP-1-mediated signaling pathways that mediate GB progression^1,3,7–8,21–22^

Our results have implications for GB therapeutic development. NRP-1 is overexpressed in GB and contributes to angiogenesis, tumor cell invasion, immune modulation, and mediates interactions with the extracellular matrix^1,3,7–8^. Current GB therapy is limited to high recurrence after surgery, radiation, and temozolomide chemotherapy, yet two-year survival rates remain low^1,3,5^.

Furthermore, the anti-VEGF antibody bevacizumab improved progression-free survival but did not improve overall survival when added to standard therapy for newly diagnosed GB^76^. This motivates for development alternative strategies that target invasion and tumor-micro environment interactions^1,3,5,7^. If peptide **1** or its optimized derivatives can bind the b1-NRP-1 pocket experimentally, these peptides could possibly interfere with VEGF-A_165_ -mediated NRP-1 signaling or other NRP-1-dependent interactions relevant to GB progression^7,9,11–12,20^. CendR peptides have C-terminal free α-carboxylate and internalize upon binding^12^. Whereas our peptides are C-terminally amidated and most likely not internalize upon binding, therefore, they can be better inhibitor NRP-1 interactions. Since peptide **1** is derived from laminin, a major ECM protein, this method of approach could provide a strategy for targeting the juncture between GB cells, vascular signaling, and extracellular matrix remodeling^21–22,29–30^.

Several limitations should be noted. First, this study is limited to empirical binding energy calculations. ΔE_b_ obtained by FoldX are conservative as well as approximate and do not completely describe the energetics of small peptide binding in a solvent-exposed pocket^38–39,72^. The single experimental method, nanoDSF, establishses an apparent interaction between the peptides and recombinant NRP-1, but thermal-shift data report global stabilization rather than site-specific engagement. The present work, therefore, serves as a rational basis for further *in silico* and *in vitro* developments rather than a demonstration of functional activity. Second, MD simulations are reliant on starting conformations, force-field parameters, simulation length, and sampling completeness^44,47,58,77^. Although multipe AF3-generated peptide-protein structures were submitted to multiple µs-cale simulations, further replicates and even more varied starting poses would bolster confidence in the observed binding behavior. Third, the present analysis focused on the isolated b1-NRP-1 domain rather than the full membrane-associated receptor environment. The nanoDSF measurements, by contrast, were performed on the complete NRP-1 ectodomain (Phe22-Lys644). The experimental and computational results were, therefore, obtained on different models, and K_D,app_ values can reflect binding anywhere on the ectodomain rather than specifically within the b1 pocket. Furthermore, pocket occupancy also varied substantially among the five trajectories for each peptide, and the FoldX estimates were derived from structural windows that were not statistically independent. Nevertheless, nanoDSF measurements established an apparent interaction between all three peptides and the recombinant ectodomain NRP-1. However, K_D,app_ do not necessarily indicate exclusive binding to the b1 domain. Competitive binding of VEGF-A_165_ with the peptides studied here would determine b1 pocket specificity^11–12,20,78^.

## 5. Conclusions

Overall, this study presents computational and experimental evidence that peptide **1**, and its analogs bind to the VEGF-A_165_ binding pocket of b1-NRP-1. During MD simulations, peptide **1** showed greater pocket occupancy and more persistent contact with key b1-NRP-1 pocket residues than either peptides **2** or **3**. FoldX binding energy estimates were comparable across all three peptides, and nanoDSF confirmed an apparent interaction with recombinant NRP-1 for all three peptides, with dissociation constants in the low-micromolar range. These findings support peptide **1** as a plausible starting point for the development of laminin-derived NRP-1 binding peptides. Although peptides **2** and **3** occupied the canonical pocket less persistently, both retained measurable binding for recombinant NRP-1, and peptide **2** showed the lowest K_D,app_ of the three peptides. Peptide **2** and **3** as anti-CendR analogs may, therefore, provide a new and distinct scaffold for development of noncanonical NRP-1 ligands and warrant further evaluation as potential inhibitors. Together these results provide a groundwork for future computational, biochemical and cell-based validations of these laminin fragment-based peptides.

### Data and Software Availability

Simulation input files, processed trajectory analysis data, representative structures, and custom scripts used for analysis and figure generation are available at Zenodo repository 10.5281/zenodo.23748004. The deposited materials include the data underlying the figures and supplementary tables. Because of large production trajectory file size, representative clustered structures will be are provided upon request from the corresponding author.

## Supporting Information

Cation-π interaction analysis, configurational-entropy plots, trajectory-specific equilibration cutoffs, occupancy of the four interaction classes, FoldX interface-frequency tables, and equilibration-cutoff sensitivity analyses.

## Acknowledgements

The work were supported by the grants of Kick for a Cure, Inc. (SL) and Creighton University School of Medicine M1 Summer Research Grant (EV). Some of the python analysis codes were developed with assistance of Claude AI (Anthropic). The authors reviewed, tested, and validated all code.

