## Supplement for "Laminin-Derived Peptides Bind Neuropilin-1 at the VEGF-A_165_ Binding Pocket"

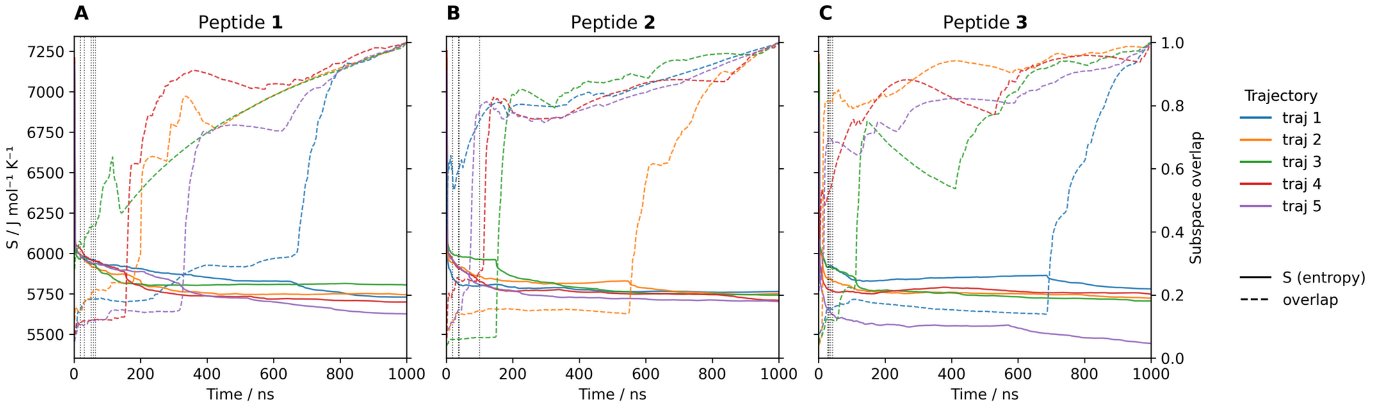


**Figure S1. Configurational-entropy traces and subspace overlap versus simulation time-step for the five independent trajectories per peptide.** A) Peptide **1** B) Peptide **2** C) Peptide **3**. In each panel, solid lines show the configurational entropy (left axis) and dashed lines the normalized subspace overlap (right axis) for the five independent trajectories, colored by trajectory. Dotted vertical lines mark each trajectory's equilibration cutoff (Supplementary Table S1); frames before the cutoff were excluded from all analyses. In every trajectory the entropy declined from ≈ 7200 J·mol$^{-1}$·K$^{-1}$ into a sustained lower plateau below ≈ 6000 J·mol$^{-1}$·K$^{-1}$, which defined the cutoff, while subspace approached 1.0, which indicates convergence of the sampled subspace.

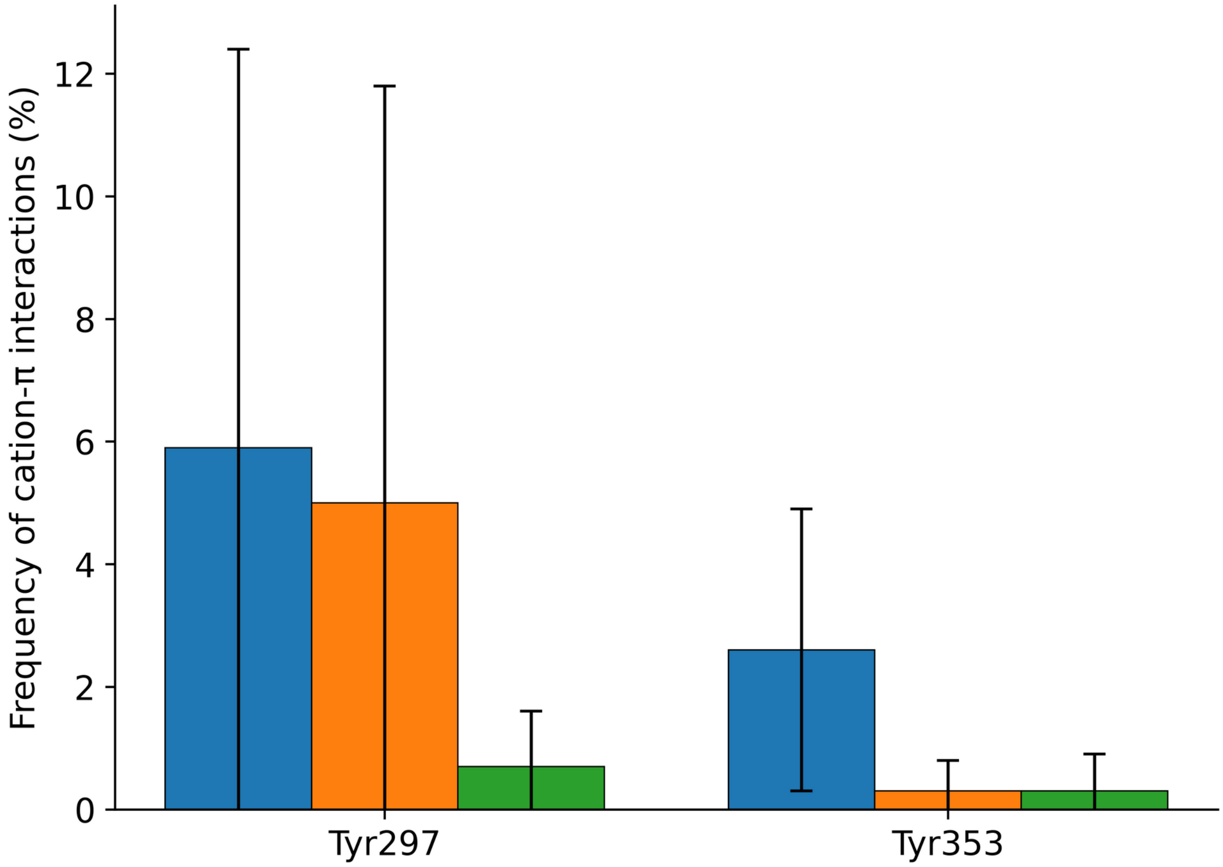


**Figure S2. Cation-π geometry between the peptide Arg guanidinium ($C_\zeta$) and the pocket Tyr297 and Tyr353.** Occupancy is the percentage of equilibrated frames satisfying a $C_\zeta$-to-ring-centroid distance ≤ 6.0 Å and an in-plane lateral offset ≤ 2.3 Å (mean ± SD across five trajectories per peptide) for peptide **1** (blue), peptide **2** (orange), and peptide **3** (green). The interaction was most frequent for peptide **1** and followed the ranking of canonical-pocket occupancy.

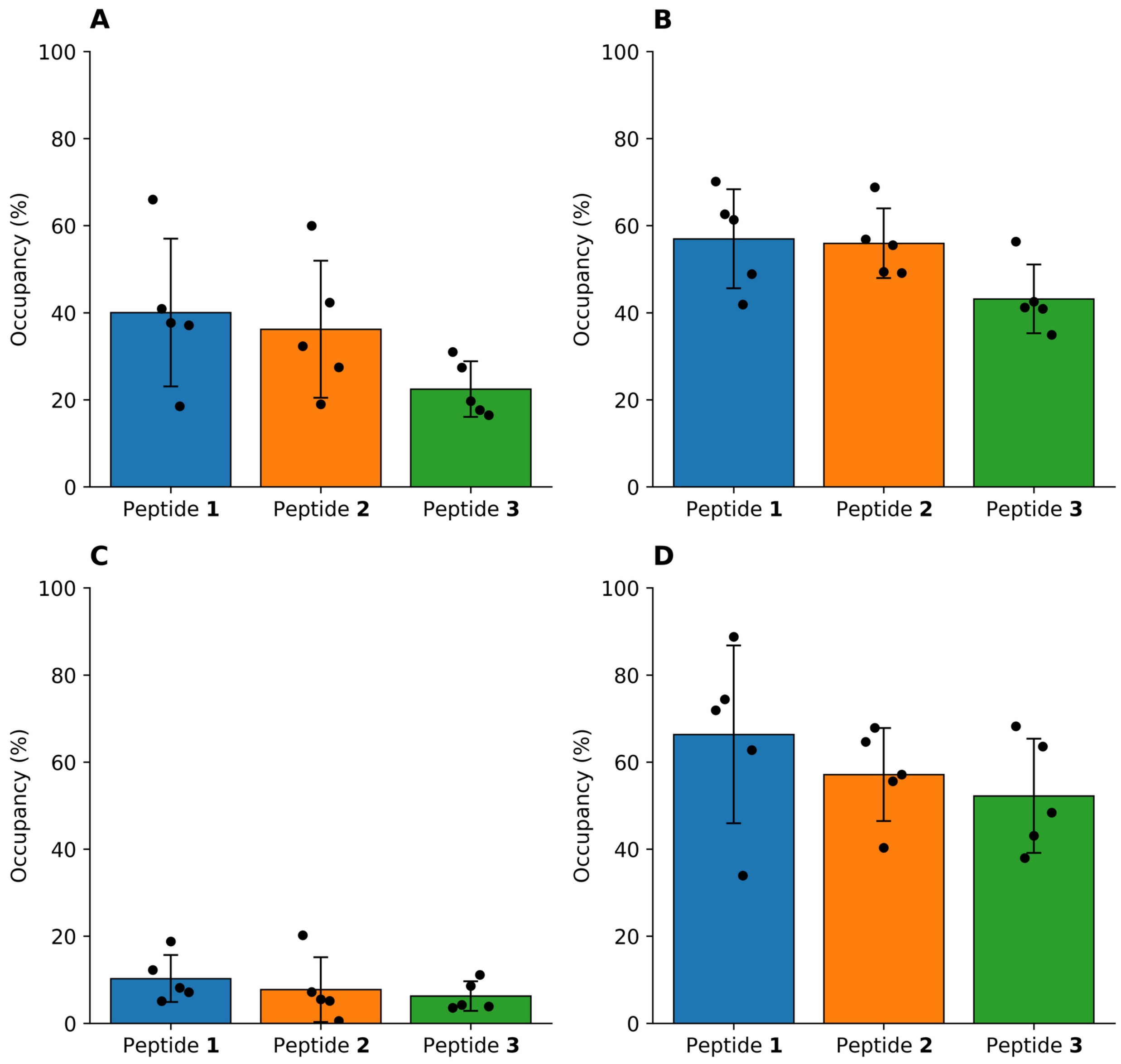


**Figure S3. Occupancy of the interaction classes over the equilibrated interval.** A) ionic/salt bridge, B) hydrogen bond, C) cation-π, D) CH-π. Bars show the mean occupancy across five independent trajectories per peptide for peptide **1** (blue), peptide **2** (orange), peptide **3** (green), error bars is the respective SD, and points the individual trajectory values. Occupancy is the percentage of equilibrated frames containing at least one interchain interaction of that class. Criteria are given in the methods.

**Table S1. Trajectory-specific post-equilibration cutoffs (ns). Values were determined independently for each of the five trajectories per peptide.**

| Trajectory | Peptide **1** | Peptide **2** | Peptide **3** |
|---|---|---|---|
| 1 | 51.0 | 37.0 | 30.9 |
| 2 | 64.3 | 19.1 | 27.7 |
| 3 | 19.1 | 36.0 | 30.1 |
| 4 | 57.6 | 38.7 | 36.3 |
| 5 | 30.9 | 99.5 | 42.4 |
| Mean ± SD | 44.6 ± 18.9 | 46.1 ± 30.9 | 33.5 ± 5.9 |

**Table S2. Per-residue FoldX interface frequency (% of equilibrated FoldX-evaluated frames; mean ± SD across trajectories) for b1-NRP-1 residues reaching ≥ 5% in at least one peptide.**

Predefined VEGF-$A_{165}$ binding pocket residues are marked with *.

| Residue | Peptide **1** | Peptide **2** | Peptide **3** |
|---|---|---|---|
| 277 | 2 ± 3 | 4 ± 7 | 10 ± 11 |
| 278 | 2 ± 3 | 8 ± 12 | 14 ± 18 |
| 282 | 2 ± 5 | 18 ± 27 | 20 ± 34 |
| 289 | 12 ± 17 | 0 ± 0 | 0 ± 0 |
| 290 | 10 ± 15 | 0 ± 0 | 0 ± 0 |
| 291 | 6 ± 13 | 4 ± 9 | 0 ± 0 |
| 292 | 13 ± 16 | 0 ± 1 | 0 ± 0 |
| 294 | 7 ± 8 | 0 ± 0 | 0 ± 1 |
| 295 | 22 ± 11 | 6 ± 8 | 14 ± 26 |
| 296 | 10 ± 8 | 4 ± 8 | 20 ± 40 |
| Tyr297* | 77 ± 29 | 37 ± 43 | 65 ± 44 |
| 298 | 16 ± 11 | 13 ± 20 | 27 ± 49 |
| 299 | 6 ± 6 | 7 ± 10 | 24 ± 47 |
| 300 | 4 ± 2 | 10 ± 13 | 8 ± 12 |
| Trp301* | 61 ± 31 | 31 ± 37 | 44 ± 43 |
| 302 | 3 ± 2 | 4 ± 6 | 17 ± 34 |
| 304 | 1 ± 1 | 4 ± 9 | 6 ± 11 |
| 308 | 3 ± 4 | 11 ± 20 | 19 ± 26 |
| 309 | 2 ± 4 | 16 ± 25 | 14 ± 25 |
| 311 | 0 ± 1 | 13 ± 22 | 0 ± 0 |
| 313 | 6 ± 4 | 8 ± 13 | 2 ± 3 |

| Residue | Peptide **1** | Peptide **2** | Peptide **3** |
|---|---|---|---|
| Thr316* | 71 ± 33 | 36 ± 42 | 47 ± 48 |
| Pro317* | 65 ± 32 | 32 ± 40 | 32 ± 38 |
| 319 | 57 ± 17 | 27 ± 34 | 28 ± 23 |
| Asp320* | 67 ± 32 | 36 ± 40 | 41 ± 42 |
| 321 | 6 ± 5 | 1 ± 1 | 5 ± 8 |
| 322 | 28 ± 30 | 13 ± 18 | 25 ± 29 |
| 323 | 3 ± 3 | 7 ± 15 | 0 ± 0 |
| 324 | 12 ± 8 | 7 ± 14 | 1 ± 1 |
| 325 | 8 ± 13 | 6 ± 14 | 0 ± 0 |
| 327 | 9 ± 16 | 0 ± 0 | 0 ± 0 |
| 328 | 6 ± 14 | 0 ± 0 | 0 ± 0 |
| 329 | 11 ± 16 | 0 ± 0 | 0 ± 0 |
| 337 | 1 ± 2 | 6 ± 8 | 28 ± 27 |
| 338 | 4 ± 6 | 19 ± 31 | 40 ± 43 |
| 342 | 0 ± 0 | 11 ± 23 | 2 ± 2 |
| 345 | 0 ± 0 | 9 ± 21 | 0 ± 0 |
| Ser346* | 34 ± 9 | 20 ± 25 | 16 ± 17 |
| 347 | 1 ± 0 | 6 ± 7 | 1 ± 1 |
| 348 | 27 ± 7 | 22 ± 25 | 20 ± 18 |
| Thr349* | 46 ± 14 | 30 ± 32 | 40 ± 43 |
| 350 | 1 ± 1 | 8 ± 10 | 2 ± 4 |
| Lys351* | 46 ± 25 | 26 ± 21 | 36 ± 43 |
| 352 | 0 ± 0 | 12 ± 17 | 0 ± 0 |
| Tyr353* | 66 ± 30 | 34 ± 37 | 44 ± 50 |
| 362 | 0 ± 0 | 6 ± 9 | 0 ± 0 |
| 364 | 1 ± 1 | 5 ± 8 | 0 ± 0 |
| 367 | 7 ± 10 | 0 ± 0 | 0 ± 0 |
| 369 | 8 ± 15 | 0 ± 0 | 0 ± 0 |
| 370 | 0 ± 0 | 6 ± 9 | 0 ± 0 |
| 374 | 0 ± 0 | 12 ± 18 | 6 ± 13 |
| 379 | 0 ± 0 | 12 ± 21 | 5 ± 10 |
| 380 | 0 ± 0 | 10 ± 19 | 1 ± 1 |
| 381 | 0 ± 0 | 12 ± 21 | 5 ± 10 |
| 382 | 0 ± 1 | 15 ± 21 | 3 ± 5 |
| 387 | 0 ± 0 | 7 ± 11 | 0 ± 0 |
| 388 | 0 ± 0 | 12 ± 23 | 0 ± 1 |
| 389 | 3 ± 5 | 15 ± 26 | 13 ± 17 |

| Residue | Peptide **1** | Peptide **2** | Peptide **3** |
|---|---|---|---|
| 390 | 4 ± 7 | 22 ± 34 | 42 ± 45 |
| 391 | 3 ± 5 | 12 ± 17 | 18 ± 20 |
| 392 | 6 ± 8 | 32 ± 44 | 52 ± 53 |
| 393 | 0 ± 0 | 14 ± 21 | 4 ± 6 |
| 394 | 4 ± 6 | 28 ± 40 | 49 ± 49 |
| 396 | 0 ± 0 | 13 ± 21 | 6 ± 12 |
| 397 | 0 ± 0 | 5 ± 7 | 1 ± 2 |
| 398 | 0 ± 0 | 5 ± 6 | 5 ± 6 |
| 399 | 0 ± 0 | 6 ± 9 | 0 ± 0 |
| 400 | 0 ± 0 | 6 ± 9 | 0 ± 0 |
| 402 | 7 ± 10 | 2 ± 3 | 0 ± 0 |
| 403 | 12 ± 17 | 0 ± 0 | 0 ± 0 |
| 407 | 0 ± 0 | 7 ± 15 | 0 ± 0 |
| 408 | 0 ± 0 | 6 ± 14 | 0 ± 0 |
| 409 | 1 ± 1 | 6 ± 14 | 0 ± 0 |
| 410 | 6 ± 7 | 5 ± 11 | 1 ± 2 |
| Trp411* | 29 ± 30 | 11 ± 18 | 19 ± 22 |
| 412 | 3 ± 4 | 8 ± 16 | 1 ± 3 |
| Thr413* | 32 ± 32 | 14 ± 13 | 34 ± 40 |
| Ile415 * | 62 ± 34 | 31 ± 32 | 35 ± 41 |
| 416 | 39 ± 35 | 11 ± 14 | 14 ± 16 |
| 418 | 0 ± 0 | 10 ± 23 | 0 ± 0 |
| 420 | 2 ± 3 | 10 ± 16 | 11 ± 13 |
| 422 | 5 ± 8 | 27 ± 36 | 48 ± 49 |

**Table S3. Sensitivity of canonical-pocket occupancy (window method, %) to the equilibrated cutoff. Pocket occupancy was recomputed with each trajectory's equilibrated start shifted 0, 50, or 100 ns later than its entropy-based cutoff.**

| Peptide | Baseline cutoff | +50 ns | +100 ns |
|---|---|---|---|
| **1** | 60.3 | 58.1 | 56.2 |
| **2** | 25.9 | 25.1 | 25.0 |
| **3** | 21.4 | 20.4 | 19.8 |

The peptide **1** > peptide **2** > peptide **3** ranking is preserved and absolute values change by no more than four percentage points, indicating the result is not sensitive to the exact cutoff.
